# Chromatin architecture in addiction circuitry elucidates biological mechanisms underlying cigarette smoking and alcohol use traits

**DOI:** 10.1101/2021.03.18.436046

**Authors:** Nancy Y.A Sey, Benxia Hu, Marina Iskhakova, Huaigu Sun, Neda Shokrian, Gabriella Ben Hutta, Jesse Marks, Bryan C. Quach, Eric O. Johnson, Dana B. Hancock, Schahram Akbarian, Hyejung Won

## Abstract

Cigarette smoking and alcohol use are among the most prevalent substances used worldwide and account for a substantial proportion of preventable morbidity and mortality, underscoring the public health significance of understanding their etiology. Genome-wide association studies (GWAS) have successfully identified genetic variants associated with cigarette smoking and alcohol use traits. However, the vast majority of risk variants reside in non-coding regions of the genome, and their target genes and neurobiological mechanisms are unknown. Chromosomal conformation mappings can address this knowledge gap by charting the interaction profiles of risk-associated regulatory variants with target genes. To investigate the functional impact of common variants associated with cigarette smoking and alcohol use traits, we applied Hi-C coupled MAGMA (H-MAGMA) built upon cortical and midbrain dopaminergic neuronal Hi-C datasets to GWAS summary statistics of nicotine dependence, cigarettes per day, problematic alcohol use, and drinks per week. The identified risk genes mapped to key pathways associated with cigarette smoking and alcohol use traits, including drug metabolic processes and neuronal apoptosis. Risk genes were highly expressed in cortical glutamatergic, midbrain dopaminergic, GABAergic, and serotonergic neurons, suggesting them as relevant cell types in understanding the mechanisms by which genetic risk factors influence cigarette smoking and alcohol use. Lastly, we identified pleiotropic genes between cigarette smoking and alcohol use traits under the assumption that they may reveal substance-agnostic, shared neurobiological mechanisms of addiction. The number of pleiotropic genes was ∼26-fold higher in dopaminergic neurons than in cortical neurons, emphasizing the critical role of ascending dopaminergic pathways in mediating general addiction phenotypes. Collectively, brain region- and neuronal subtype-specific 3D genome architecture refines neurobiological hypotheses for smoking, alcohol, and general addiction phenotypes by linking genetic risk factors to their target genes.

## Introduction

The National Survey on Drug Use and Health in 2018 estimated that 27.3 million individuals were daily cigarette smokers and 16.6 million individuals were heavy alcohol users^1^. Cigarette smoking and alcohol use are the 1^st^ and 3^rd^ leading causes of mortality and morbidity, accounting for 480,000 and 88,000 deaths per year in the United States, respectively^2, 3^. Despite their public health burden, treatment options for nicotine and alcohol use disorders are limited and often fail. However, existing treatments can be improved and new treatments can be developed with a better understanding of the underlying neurobiology of addiction. Genome wide association studies (GWAS) on smoking and alcohol use traits have demonstrated that common variation explains a significant proportion of phenotypic variance of substance use^4^. Nearly 400 genomic loci were found to have an impact on smoking and/or alcohol use traits from GWAS sample sizes up to 1.2 million^4–7^. However, the vast majority of associated variants reside in non-coding DNA, and their target genes and relevant neurobiological mechanisms are poorly understood. Examining higher-order chromatin architecture is crucial to understanding the functional consequences of non-coding variation by linking variants to distal genes based on chromatin interaction profiles^8–10^. Whereas the three-dimensional (3D) genomic landscape of the human brain has advanced our understanding of neurobiological mechanisms underlying psychiatric disorders^9, 11–13^, such approaches have been essentially lacking in explaining the genetic architecture of substance use disorders (SUD).

To understand the functional impact of common variants associated with cigarette smoking and alcohol use, we applied Hi-C coupled MAGMA (H-MAGMA)^12^ to GWAS of smoking and alcohol use traits and identified their putative target genes for further characterization^5–7^. Smoking and alcohol use traits likely affect neurocircuits that underlie addiction and include the prefrontal cortex (PFC), nucleus accumbens (NAc), amygdala, and midbrain dopaminergic cell groups such as ventral tegmental area (VTA) and substantia nigra (SN)^14–16^. We reasoned that characterization of chromatin architecture across the brain reward circuitry is critical to understanding the gene regulatory mechanisms associated with substance use. With neurons being the major drivers of substance use behaviors, we profiled chromatin architecture from cortical neurons (CNs) in the dorsolateral PFC (DLPFC)^17^ and dopaminergic neurons (DNs) in the midbrain^18^. We then built H-MAGMA inputs from CNs and DNs, and applied them to GWAS summary statistics of smoking and alcohol use traits. In particular, given the recent work on a potential difference in genetic architecture between substance consumption and clinical diagnosis of use disorder^4^, we mapped genetic variants associated with consumption or use (drinks per week [DPW^5^] and cigarettes per week [CPD])^5^ versus use disorder (problematic alcohol use [PAU]^6^ and nicotine dependence [ND]^7^) to their associated risk genes. Our analysis of substance use risk genes identified key biological pathways, primary cell types, and brain circuitry that might confer risk for substance use. In addition, we characterized genes and pathways shared between cigarette smoking and alcohol use traits to provide a core neurobiological basis of addiction.

## Result

### Epigenetic landscape of cortical and midbrain dopaminergic neurons

Neural circuitry underlying addiction involves, among others, dopaminergic cell groups in the midbrain, including VTA and SN, as well as neuronal populations in the PFC^14^ (**Figure 1A**). However, the gene regulatory landscape in these two brain regions and its implication in genetics of cigarette smoking and alcohol use traits have not been studied. To understand the relationship between the reward circuitry and genetic underpinnings of substance use, we evaluated enrichment of genetic risk factors for four traits associated with alcohol use (PAU^6^ and DPW^5^) and cigarette smoking (ND^7^ and CPD^5^) in *cis*-regulatory elements (CREs) of the midbrain and PFC^19^ (**Methods**). Using stratified LDSC, we demonstrated that every trait showed significant heritability enrichment for CREs in the midbrain and PFC (**Figure 1B**).

**Figure 1.**
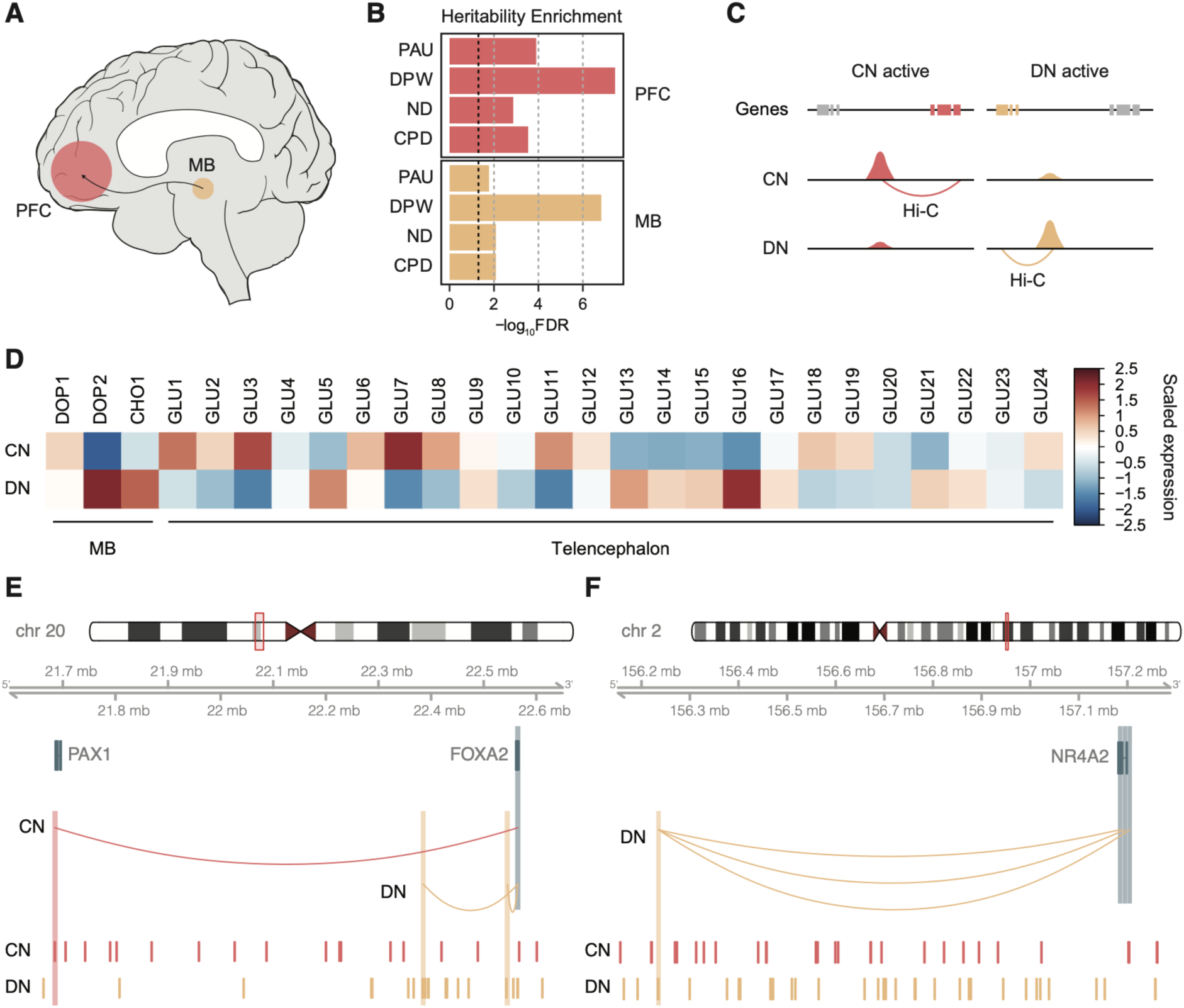
Gene regulatory landscape in cortical and dopaminergic neurons. **A.** Brain reward circuitry encompasses the midbrain (MB) and its projection to the prefrontal cortex (PFC). **B.** *Cis-* regulatory elements (CREs) in the PFC and substantia nigra (SN) are enriched for genetic risk factors for problematic alcohol use (PAU), drinks per week (DPW), nicotine dependence (ND), and cigarettes per day (CPD). The black dotted line represents *FDR=0.05*. **C.** Dopaminergic neuronal (DN) CREs were linked to their target genes using DN Hi-C data, while cortical neuronal (CN) CREs were linked to target genes using CN Hi-C data. **D.** Genes mapped to DN-CREs were highly expressed in midbrain dopaminergic (DOP2) and cholinergic neurons (CHO1), while genes mapped to CN-CREs were highly expressed in telencephalic glutamatergic neurons (GLU1, 3, 7, 11). **E-F.** Different enhancer connectivity between CNs and DNs for *FOXA2* (**E**) and *NR4A2* (**F**) loci. Promoters of *FOXA2* and *NR4A2* are highlighted in blue, while their interaction targets in CN and DN are highlighted in red and orange, respectively.

Midbrain DNs have long been hypothesized to be the major player of the brain reward circuitry^14, 20^. Thus, we investigated whether adult midbrain DN-CREs explained the heritability enrichment of cigarette smoking and alcohol use traits (**Methods**). Indeed, genetic risk factors for substance use traits were enriched in chromatin accessible regions of DNs derived from human induced pluripotent stem cells^21^ (hiPSC, **Supplementary Figure 1A**).

Given the cellular heterogeneity of the PFC, we also evaluated heritability enrichment of substance use traits in CREs of four major cell types (neurons, astrocytes, microglia, and oligodendrocytes) in the cortex^22^. Neurons showed the strongest heritability enrichment of substance use traits among the four cell types (**Supplementary Figure 1B**). These results collectively suggest that the gene regulatory relationships in CNs and DNs may provide rich information about genetic underpinnings of substance use traits.

We next sought to compare gene regulatory relationships between CNs and DNs. Whereas substantial differences in chromatin architecture have been observed across different cell types in the human brain^13, 17, 22^, little information is available for the chromatin architecture in different brain regions and/or neuronal subtypes. We linked differential chromatin accessibility peaks between CNs and DNs^21^ to their target genes on the basis of corresponding chromatin loops (**Methods**, **Figure 1C**). We then measured cell-type specific expression profiles of the genes linked to CN- and DN-CREs. Genes linked to CN-CREs were highly expressed in cortical pyramidal neurons of the telencephalon (GLU1-3, 6-8), whereas genes linked to DN-CREs were highly expressed in midbrain dopaminergic (DOP2) and cholinergic neurons (CHO1) as well as subcortical-projecting glutamatergic neurons in the telencephalon (GLU5, 13-17)^23^ (**Figure 1D**).

We also found evidence of different enhancer wiring of dopaminergic marker genes between CNs and DNs. For example, *FOXA2* and *NR4A2*, master regulators for dopaminergic neuronal specification and differentiation^24–26^, displayed different regulatory connections between CNs and DNs. *FOXA2* was linked to two proximal enhancers in DNs as compared to one distal enhancer in CNs (**Figure 1E**). In contrast, *NR4A2* was linked to multiple distal enhancers only in DNs, but not in CNs (**Figure 1F**).

We next compared topologically associating domains (TADs) between CNs and DNs (**Methods**). Consistent with previous reports^27^, TADs were largely conserved between CNs and DNs. However, we noted some differences in TAD boundary strengths (defined by *binSignal*, see **Methods** for details) between CNs and DNs. For example, *EN1 and EN2* are critical survival factors for DN differentiation and maintenance^28^. We found that *EN1* was located at the TAD boundary whose strength is stronger in DNs than in CNs (**Supplementary Figure 2A**). In contrast, a large DN TAD in which *EN2* is located was partitioned into two TADs in CNs (**Supplementary Figure 2B**). *FOXA2* also showed strengthened TAD boundaries in DNs (**Supplementary Figure 2C**), which corresponds to the confinement of loops in proximal space as evidenced in **Figure 1E**. Importantly, these genes were more highly expressed in DNs than in CNs (**Supplementary Figure 2**). Therefore, these results indicate that different neuronal subtypes involved in substance use traits display distinct chromatin architecture that is coupled with transcriptional regulation.

### CN and DN H-MAGMA identifies genes and biological pathways underlying cigarette smoking and alcohol use traits

To investigate the functional impact of common variants associated with cigarette smoking and alcohol use traits, we next employed H-MAGMA to assign genetic variants to their target genes based on long-range chromatin interaction^12^. As heritability enrichment results suggested the role of CNs and DNs in cigarette smoking and alcohol use traits (**Figure 1B, Supplementary Figure 1**), we generated H-MAGMA input files from CN and DN Hi-C data (hereafter referred to as CN and DN H-MAGMA, respectively). We applied H-MAGMA to PAU, DPW, ND, and CPD, and identified risk genes for each trait using a false discovery rate (*FDR*) threshold of 5% (**Figure 2A-B, Supplementary Tables 2 and 3**). We detected a small number of risk genes for ND in comparison to other GWAS, which can be attributed to the smaller sample size of ND GWAS (see **Methods** for the sample size for each GWAS).

**Figure 2.**
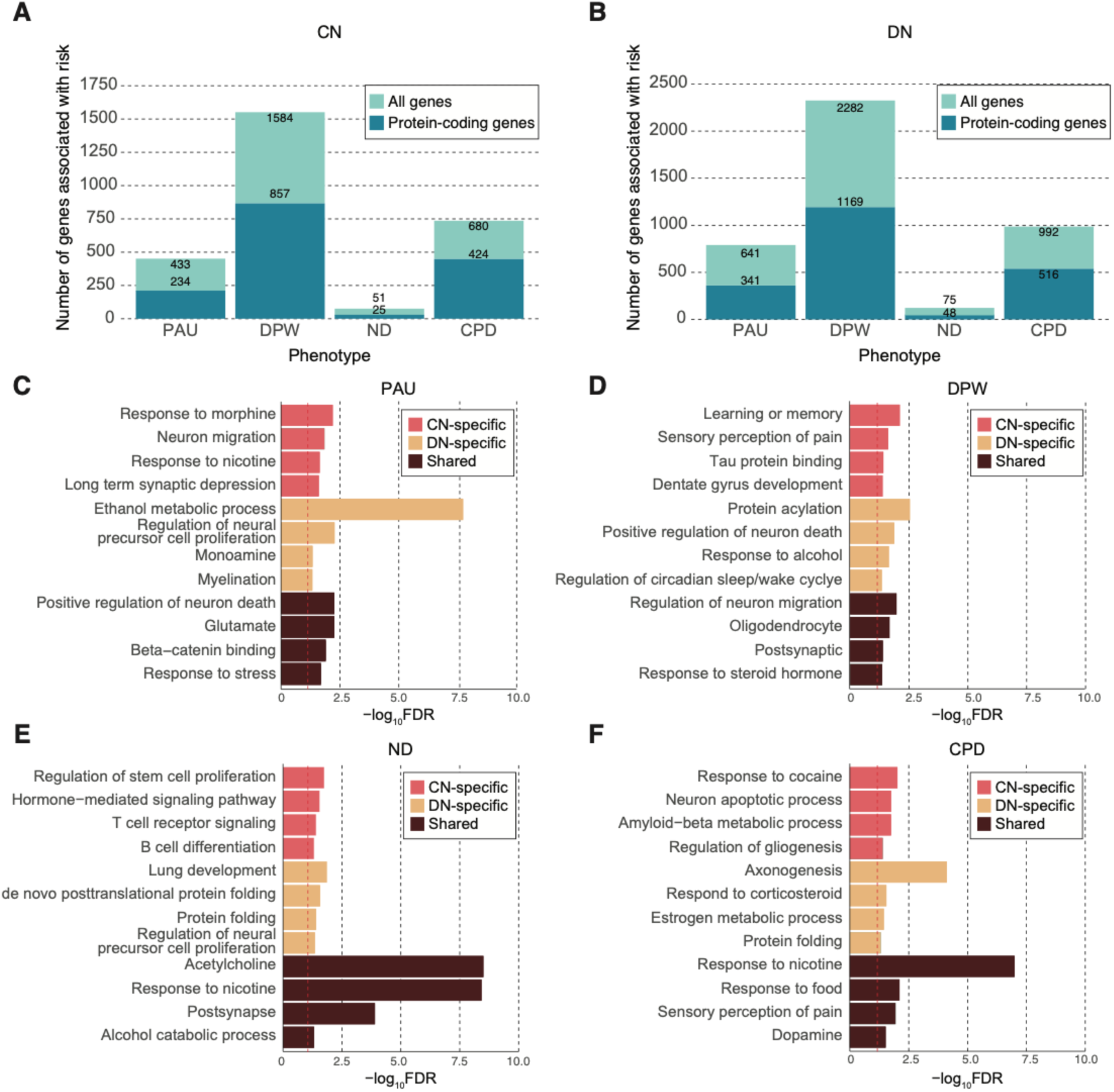
Genes and pathways associated with cigarette smoking and alcohol use traits. **A-B.** The number of risk genes for cigarette smoking and alcohol use traits based on H-MAGMA built from CN **(A)** and DN **(B)** Hi-C data (*FDR<0.05*). For each stacked bar plot, an upper bar plot in light blue denotes all genes, whereas a lower layer in dark blue corresponds to protein-coding genes. **C-F**. Gene ontologies (GO) enriched for PAU **(C)**, DPW **(D)**, ND **(E)**, and CPD **(F)**. CN-specific GO terms represent terms unique to genes identified from H-MAGMA built on CN Hi-C data, while DN-specific GO terms represent terms unique to genes identified from H-MAGMA built from DN Hi-C data. Shared terms denote GO terms detected in both CN and DN H-MAGMA results. Dotted red line denotes *FDR=0.05*.

Next, we mapped risk genes identified from CN and DN H-MAGMA to biological pathways using gene ontology (GO) analysis. Rather than using a specific FDR threshold, we ran ranked-based GO analysis using the Z-score of H-MAGMA output files (**Methods**). Since we used two separate H-MAGMA inputs to assign common variants to their target genes, we obtained two GO results for each trait – one for CN H-MAGMA risk genes and the other for DN H-MAGMA risk genes. We then classified the results as CN-specific GO terms to represent biological pathways unique to CN H-MAGMA. Comparably, we classified DN-specific GO terms to represent biological pathways unique to DN H-MAGMA.

We validated previous findings that ethanol metabolic processes and response to alcohol were associated with PAU and DPW (**Figure 2C-D**)^4, 6^, and that cholinergic and nicotinic pathways were associated with ND and CPD^5, 7^ (**Figure 2E-F**, see **Supplementary Table 4** for full GO outputs). Notably, we also identified alcohol catabolic processes for ND and nicotinic pathways for PAU.

Likewise, we further identified GO terms relating to other substances of abuse. For instance, GO terms for PAU included response to morphine (**Figure 2C**), while GO terms for CPD included response to cocaine (**Figure 2F**). Taken together, these findings underscore potential genetic overlap and interplay among different substances of abuse.

We identified several similarities across cigarette smoking and alcohol use traits. For example, neuronal processes such as neuronal migration and apoptosis were associated with cigarette smoking and alcohol use, which is in line with studies that have pinpointed the disruption of neuronal migration and neurotransmission in response to substance use^29–31^. We also observed myelination and gliogenesis to be associated with DPW and CPD, respectively, hinting at the role of neuron-glia interactions in substance use traits^32, 33^. Several immune processes including T and B cell activation were shown to be associated with cigarette smoking and alcohol use, which corroborates the relationship between substance use and suppressed immunity (**Figure 2E**)^34–38^. We also identified a potential role of protein folding that has been shown to contribute to the stress response^39^. A potential link between substance use and neurodegeneration emerged, such as amyloid-beta metabolic processes for CPD and tau protein binding for DPW^40, 41^. Lastly, pain perception was associated with DPW and CPD, consistent with prior research linking pain perception and the reward circuitry^42, 43^.

We also observed distinct biological processes between cigarette smoking and alcohol use traits. For instance, long term synaptic depression (**Figure 2C**) as well as learning and memory (**Figure 2D**) were characteristic of alcohol use traits but not cigarette smoking, highlighting the important role of synaptic plasticity and memory consolidation in the mechanism of alcohol use^44, 45^. We also found GO terms relating to sleep and wake cycle for alcohol use traits, which support a rich body of evidence suggesting that prolonged alcohol use and misuse can cause deleterious effects on sleep quality^46, 47^. Cigarette smoking traits also exhibited distinct associations not observed in alcohol use traits. For instance, we noted lung development to be associated with both ND and CPD which supports epidemiological findings of lung morbidities linked to cigarette smoking^48, 49^.

Discrete biological processes were also observed between CN and DN H-MAGMA. Ethanol metabolism and alcohol response were enriched for alcohol use traits in a DN-specific manner (**Figure 2C-D**). In contrast, the potential link between neurodegeneration and substance use was specific to CNs. These results suggest that the neurobiological basis of cigarette smoking and alcohol use traits may need to be studied in a brain region- and neuronal subtype-specific manner.

### Cellular expression profiles of cigarette smoking and alcohol use risk genes convey cell types associated with substance use

Since CNs and DNs display heterogeneity and act in synchrony with multiple cell types, we leveraged single-cell RNA sequencing (scRNA-seq) datasets to further refine neuronal subtypes that confer risk of substance use. We first evaluated cellular expression profiles of cigarette smoking and alcohol use risk genes identified from CN H-MAGMA using scRNA-seq data from the human cortex^50^. We not only recapitulated our findings that genetic risk variants underlying cigarette smoking and alcohol use are enriched for neurons, but also observed that the risk genes were highly expressed in excitatory neurons (**Figure 3A**). Specifically, we found PAU, DPW, and CPD to be enriched for layer 5 pyramidal neurons (Ex5) that project to both cortical and subcortical areas including the striatum and the midbrain^51, 52^ and layer 4 neurons (Ex2) that receive sensory signals from the thalamus, a region that has been shown to be integral to addiction by modulating arousal and motivation^53^.

**Figure 3.**
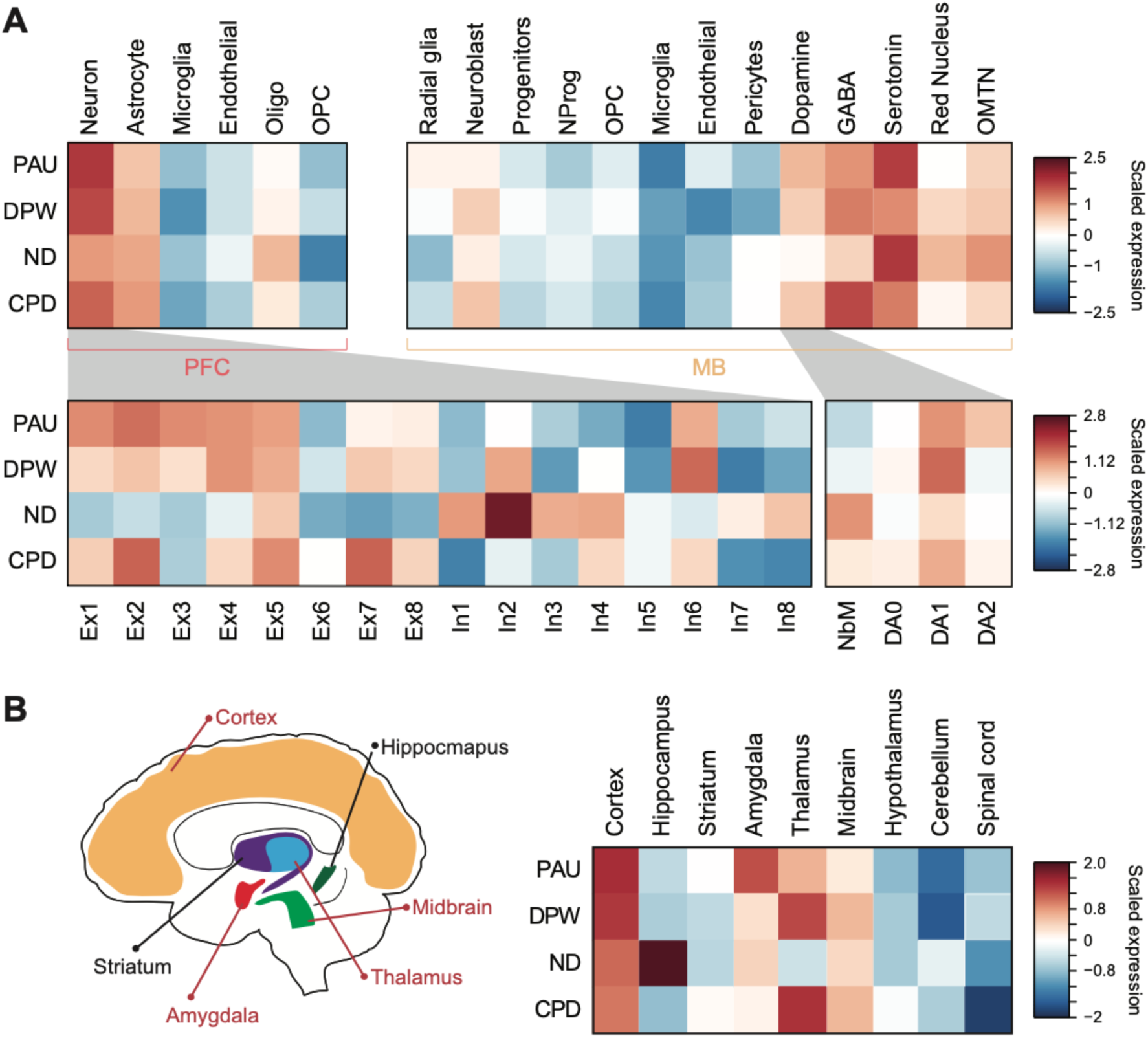
Cellular and brain regional expression profiles of cigarette smoking and alcohol use risk genes. **A.** Top left panel represents cellular expression profiles of cigarette smoking and alcohol use risk genes identified from CN H-MAGMA using scRNA-seq data from the adult cortex^50, 62^. Genetic risk factors underlying cigarette smoking and alcohol use exhibit strong enrichment in neurons. Bottom left panel represents risk gene expression across neuronal subclusters. OPC, oligodendrocytes progenitor cells; Ex, excitatory neurons; In, inhibitory neurons. Top right panel, cellular expression profiles of cigarette smoking and alcohol use risk genes identified from DN H-MAGMA using scRNA-seq from the ventral midbrain of human embryo^54^. We identified enrichment for dopaminergic, GABA-ergic, and serotonergic neurons in the midbrain. NProg, neuronal progenitors; OMNT, oculomotor and trochlear nucleus. Bottom right panel, cigarette smoking and alcohol use risk genes were enriched for DA1 across DN development in human embryonic midbrain. NbM, medial neuroblasts and precursors of DNs; DA0, immature DNs; DA1, intermediate DNs; DA2 matured DNs. **B.** Left, graphic representation of brain regions with elevated expression levels of risk genes for substance use traits. Regions highlighted in red are enriched for at least three of the four traits. Right, brain regional expression profiles of cigarette smoking and alcohol use risk genes using scRNA-seq from the mouse nervous system^23^. We detected enrichment spanning multiple brain regions including cortex, amygdala, and midbrain.

Comparably, we examined expression profiles of cigarette smoking and alcohol use risk genes identified from DN H-MAGMA in midbrain cell types using scRNA-seq data from the human embryonic ventral midbrain^54^. DNs were enriched for all traits except for ND, providing additional evidence to support the impact of DNs in modulating substance use via the reward-circuitry^14^ (**Figure 3A**). Within the DN lineage, we found enrichment for intermediate DNs (DA1) for all traits, suggesting that they may be more vulnerable to substance use. Moreover, we found enrichment for midbrain GABAergic neurons which have been shown to regulate a diverse set of processes including motor control and inhibition of dopaminergic cells, thereby modulating the reward-circuitry^55, 56^. Similarly, the observed enrichment of serotonergic neurons is consistent with their reported involvement in substance use vulnerability^57^.

Next, we extended our approach to a brain-wide fashion by assessing brain regional expression profiles of cigarette smoking and alcohol use risk genes. We leveraged extensive scRNA-seq data from the mouse nervous system to determine brain regions with high expression values of risk genes identified by CN and DN H-MAGMA^23^. Both cigarette smoking and alcohol use risk genes were highly expressed in cortical and midbrain regions as expected (**Figure 3B**). We also found strong enrichment in the hippocampus for ND risk genes, highlighting the role of hippocampus-dependent learning in ND^58, 59^. Furthermore, thalamic expression was observed for PAU, DPW, and CPD risk genes, which is consistent with the enrichment for Ex2 that receives thalamic inputs (**Figure 3A**) and points to the role of sensory perception in drug seeking behaviors^60, 61^. Finally, our results highlight enrichment of amygdala for risk genes associated with cigarette smoking and alcohol use. The association of risk variants with amygdala underscores the role of emotional processing in substance use due to its projections to other parts of the reward-circuitry^14^.

### Shared genetic architecture among substance use

Individuals often become dependent on multiple substances, and these comorbidities may be driven by shared genetic signal^6, 63^. We hypothesized that biological characterization of pleiotropic genes between cigarette smoking and alcohol use traits would identify neurobiological mechanisms underlying shared genetic architecture of substance use traits.

We first calculated genetic correlations and gene-level overlap across cigarette smoking and alcohol use traits using LD score regression (LDSC)^64^ and rank-rank hypergeometric overlap (RRHO) test^65^, respectively (**Supplementary Figure 3**). We found that RRHO of DN H-MAGMA outputs gives stronger gene-level overlaps than that of CN H-MAGMA. For example, 119 and 3,120 genes were shared between PAU and CPD using CN and DN H-MAGMA, respectively (**Figure 4A**, **Supplementary Figure 3C**). These results suggest that DNs may play a central role in explaining comorbidity in substance use. Because the PAU and CPD showed a significant genetic correlation (genetic correlation = 0.19) and gene-level overlap (RRHO Z-score = 12.16), we selected shared genes between PAU and CPD in DN H-MAGMA to serve as pleiotropic genes (**Figure 4A, Supplementary Table 5**). Pleiotropic genes were enriched for synaptic function and cell junction organization (**Figure 4B**), suggesting that alterations in synaptic organization may influence core features of substance use. We further evaluated cellular expression profiles of pleiotropic genes in the human embryonic ventral midbrain. We again found enrichment for dopaminergic, GABAergic, and serotonergic neurons, indicating their potential function in substance use biology (**Figure 4C**).

**Figure 4.**
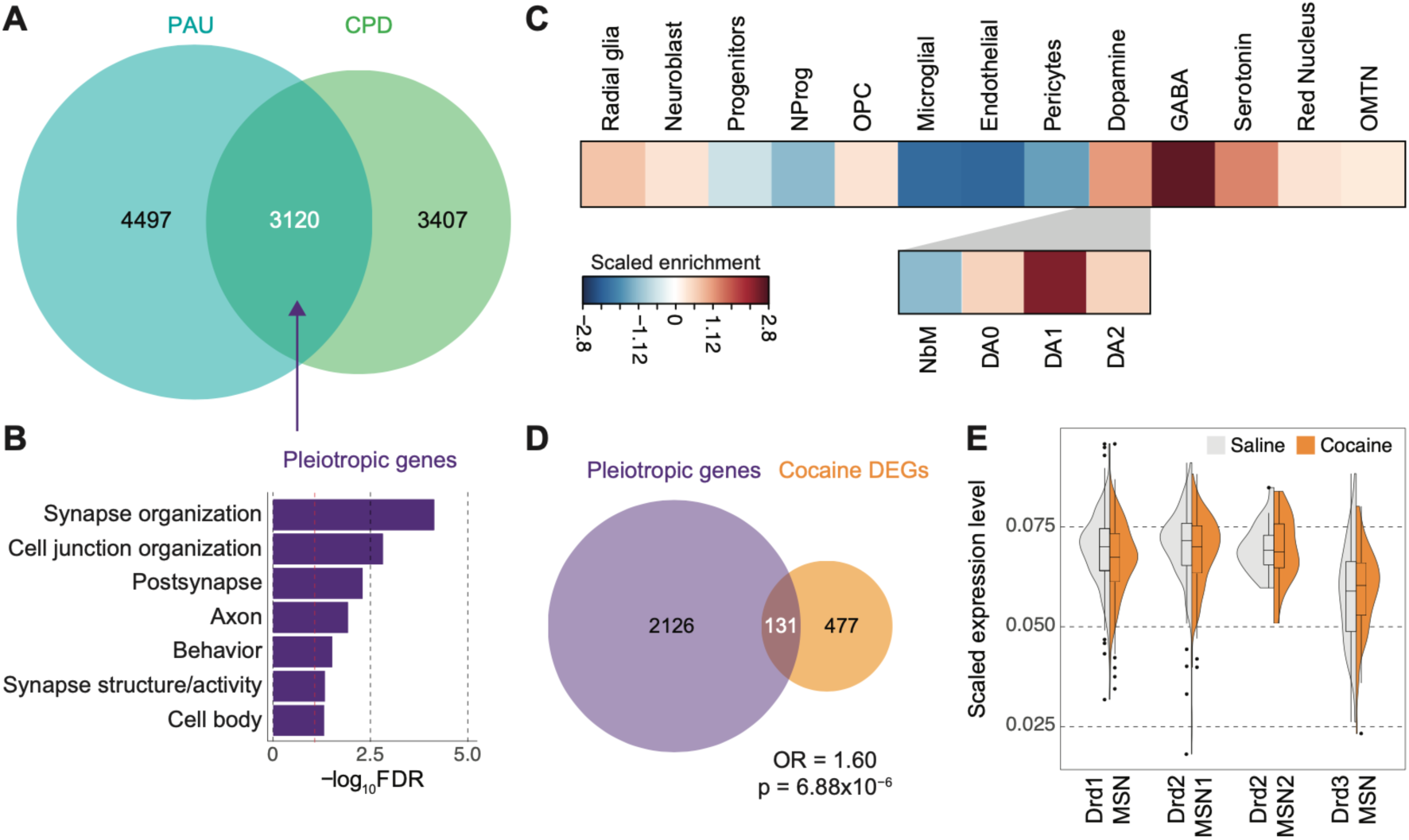
Pleiotropic genes highlight shared neurobiological bases of cigarette smoking and alcohol use. **A.** Overlap between PAU and CPD risk genes identified by DN H-MAGMA using a rank rank hypergeometric overlap (RRHO) test. Overlapping genes represent pleiotropic genes. **B**. Biological processes and molecular functions enriched for pleiotropic genes. Dotted red line denotes *FDR=0.05*. **C.** Cellular expression profiles of pleiotropic genes in the midbrain (top plot) and dopaminergic lineage (bottom plot). Pleiotropic genes are enriched for GABAergic midbrain neurons and intermediate DNs (DA1). **D.** Overlap between pleiotropic genes and differentially expressed genes (DEGs) in the mouse NAc after cocaine treatment (Fisher’s exact test, *p*=6.88×10^-6^; odds ratio [OR]=1.60; 95% confidence interval [CI]=1.34-1.96). **E.** Cellular expression changes of pleiotropic genes in response to cocaine treatment^66^. The x-axis indicates dopaminergic clusters identified in the mouse NAc while the y-axis indicates scaled expression values of the pleiotropic genes in each cluster. Drd1-MSN, dopamine receptor D1 medium spiny neurons enriched for D1-like family; Drd2-MSN and Drd2-MSN2, dopamine receptor D2 medium spiny neurons; Drd3-MSN, dopamine receptor D3 medium spiny neurons.

Based on our hypothesis that pleiotropic genes between cigarette smoking and alcohol use traits may represent risk genes shared across multiple SUD, we next examined whether they are dysregulated in response to other substances. We overlapped our pleiotropic genes with differentially expressed genes (DEGs) in the mouse NAc after cocaine treatment^66^. We found a significant proportion of our pleiotropic genes was dysregulated in response to cocaine (**Figure 4D**). We also compared the cellular expression profiles of pleiotropic genes in a saline versus cocaine treatment condition^66^. We found that pleiotropic genes were downregulated in response to cocaine in clusters of DNs that represent D1- and D2-type medium spiny neurons (**Figure 4E, Supplementary Figure 4**). Taken together, these results indicate that pleiotropic genes derived from cigarette smoking and alcohol use traits can provide insights into the core neurobiological mechanism of substance abuse.

### Drug repurposing analysis

A fundamental issue facing the treatment of SUD is the limited number of effective medications available. Although medications such as Naltrexone^67^ and Nicotine Replacement Therapies (NRT)^68^ have been traditionally used to treat alcohol use disorder and nicotine addiction, respectively, their efficacies are lacking or produce severe adverse outcomes, rendering the need for new treatment. To address this challenge, we used the Drug Signature and Drug Matrix databases of EnrichR^69^, a comprehensive gene analysis tool to identify potential drug candidates for SUD based on genetic evidence. We identified several significantly enriched drug candidates for cigarette smoking and alcohol use traits (**Supplementary Table 6**). Among these included mood stabilizers and selective serotonin reuptake inhibitors such as Fluoxetine, Citalopram, and Imipramine, consistent with their potential therapeutic benefits in some patients diagnosed with nicotine or alcohol dependency^72, 73^. We further identified enrichment for antipsychotics such as Chlorpromazine and Clozapine, pointing to some degree of convergence of addiction-relevant risk genes with molecular pathways implicated in other types of psychiatric illnesses. These findings speak to the well-documented epidemiological^70, 71^ and genetic^72, 73^ evidence supporting the comorbidity between psychiatric illnesses and substance use.

## Discussion

We interrogated chromatin interaction profiles of CNs and DNs, two primary neuronal subtypes involved in the neurocircuitry of addiction, to map GWAS risk variants of cigarette smoking and alcohol use traits to their target genes.

We built enhancer-promoter interaction landscapes in CNs and DNs by combining Hi-C and ATAC-seq, and demonstrated brain region- and neuronal subtype-specific gene regulatory relationships. We then employed these profiles to perform CN and DN H-MAGMA, which was used to identify risk genes and neurobiological pathways underlying PAU, DPW, ND, and CPD. Investigation into the biological pathways underlying cigarette smoking and alcohol use risk genes revealed the important role of drug catabolic process and alcohol metabolic process in substance use. Notably, we found that substance use risk genes were enriched for pathways associated with other neurodegenerative disorders such as tau protein binding for DPW and amyloid-beta metabolic process for CPD. The association between substance use and neurodegenerative disorders has been observed in a mouse model of Alzheimer’s disease where alcohol exposure was shown to heighten neuronal and behavioral deficits related to Alzheimer’s disease^74^. Thus, our results provide additional evidence to support that substance use and neurodegenerative disorders may share underlying genetic risk factors^75, 76^ and that risk variants associated with alcohol use may exacerbate neurodegenerative disorders by disrupting protein metabolism. We also identified an association between cigarette smoking and food intake which is in line with the previous reports linking weight gain with smoking cessation^77, 78^.

We next surveyed the cellular expression profiles of cigarette smoking and alcohol use risk genes to refine cortical and midbrain neuronal subtypes that confer risk for substance use. Within CNs, we found that cigarette smoking and alcohol use risk genes were highly expressed in glutamatergic neurons, providing an additional level of support for the neuronal basis of addiction^14^. Interestingly, we have previously shown that risk genes of psychiatric disorders were also enriched for glutamatergic neurons^12^. Based on prior epidemiological studies reporting higher substance use among individuals with mental health issues, these results suggest potential cellular basis of comorbidity between substance use and psychiatric disorders^63^. We also identified potential divergence between ND and CPD such that, risk genes associated with ND were enriched for inhibitory neurons, in contrast to the observed excitatory enrichment for CPD risk genes. While this may hint to distinct biological patterns underlying use (CPD) versus a use disorder (ND), caution should be exercised as this finding could also be influenced by the smaller number of genes associated with ND in comparison to CPD due to the smaller sample size of ND GWAS. Our cellular expression profiles within the DN lineage showed enrichment for intermediate DNs (DA1), suggesting early development as a critical time period for risk of substance use^79, 80^. Finally, we leveraged cigarette smoking and alcohol use risk genes to identify brain circuitry of addiction based on the hypothesis that defining brain regions most relevant to substance use may help derive better targeted approaches to treating SUD. In addition to the cortical and midbrain enrichment, we found enrichment for amygdala and thalamus, reinforcing that multiple brain regions are important for understanding substance use and addiction.

To further characterize how cigarette smoking and alcohol use risk genes can expand our understanding of substance use and addiction as a whole, we generated a list of pleiotropic genes between PAU and CPD. In contrast to individual risk genes being more focused on individual substance use traits, we reasoned that pleiotropic genes would provide us with the opportunity to identify principal pathways associated with addiction. Therefore, we generated pleiotropic genes using both CN and DN H-MAGMA output files. DNs, but not CNs, showed strong gene-level overlap between PAU and CPD, conveying that DNs might be the central cell type that mediates pleiotropy. Based on our hypothesis that pleiotropic genes might translate beyond just cigarette smoking and alcohol use traits, we compared them with DEGs in response to cocaine^66^. Indeed, we showed that pleiotropic genes were likely to be dysregulated in response to cocaine in the mouse NAc, demonstrating that these genes may be more susceptible to a wide range of substance use.

Lastly, we took advantage of EnrichR to identify potential drug candidates to treat ND and alcohol use disorder. We found potential drug candidates including those already on the market to treat various psychiatric illnesses such as depression and schizophrenia, further supporting a shared genetic architecture between psychiatric illnesses and substance use. Together, we demonstrate that H-MAGMA built from brain region- and neuronal subtype-specific chromatin architecture can successfully identify risk genes and biologically relevant processes associated with cigarette smoking and alcohol use.

## Methods

### Nuclei sorting

Brain tissue was cut and dounced with 5 mL of lysis buffer with RNase inhibitor, then transferred to an ultracentrifuge tube, followed by immediately adding 9 mL of sucrose buffer underlaid beneath the solution (see **Supplementary Table 1** for sample information). The samples were then spun at 24,000 rpm in an ultracentrifuge for 1 hour at 4°C. Next, the pellet was resuspended with 1mL of 0.1% BSA in DBPS, which was subsequently left on ice for 5-10 minutes. Pre-conjugated Nurr1 primary antibody (N4664) that had been incubated with the secondary antibody (Alexa 647) for an hour was then added to the nuclei suspension. Subsequently, 1.5 uL of NeuN antibody conjugated with Alexa 488 was added. Samples were wrapped in foil and rotated for 2 hours at 4°C. After 2 hours of incubation, DAPI was added to the reaction. The nuclei suspension is immediately taken to be processed on a FACSAria flow cytometry sorter, with all gates modified to eliminate debris and divide cells effectively, resulting in an apparent separation of nuclei populations through their fluorescent cell signal.

### Dopaminergic neuronal Hi-C library generation

Post-FACS sorting, 6,000 dopaminergic neuronal nuclei (NeuN+/Nurr1+) were processed through the Arima-HiC Kit User Guide for Mammalian Cell Lines (A51008, San Diego, CA) according to the manufacturer’s instructions. Afterwards, genomic DNA was purified using the Beckman Coulter AMPure® SPRIselect Beads (Indianapolis, IN). Subsequently, samples were sonicated utilizing the Covaris S220 (Woburn, MA), then size selected and purified using Beckman Coulter AMPure® SPRIselect Beads (Indianapolis, IN) to target for 300–500 base pair sized fragments. Samples were then enriched in biotin using the Arima-HiC Kit for Library Preparation, alongside the Swift Biosciences® Accel-NGS® 2S Plus DNA Library Kit (San Diego, CA). Afterwards, the Swift Biosciences Accel-NGS 2S Plus DNA library kit (21024, Ann Arbor, MI) was utilized for end repair and adapter ligation. Unique indices were ligated to each sample using the Swift Biosciences 2S Indexing Kit (26148). DNA libraries were amplified and purified using the Kapa Hyper Prep Kit (NC0709851, Wilmington, MA) and Beckman Coulter AMPure® SPRIselect Beads according to the manufacturer’s instructions. Resulting Hi-C libraries were sequenced through Illumina NovaSeq 6000 (150 bp paired-end sequencing) at a depth of 400 million reads per sample.

### Hi-C analysis

We applied HiC-Pro (v2.11.1)^81^ to the DN Hi-C^18^. In brief, we used Bowtie2 (v2.3.5.1)^82^ with *--very-sensitive -L 30 --score-min L,-0.6,-0.2 --end-to-end --reorder* to align Hi-C reads to hg19 from UCSC database, and obtained unique mapped read pairs (valid pairs). Valid pairs were then used to generate Hi-C contact matrices at 10kb and 40kb resolutions. Hi-C contact matrices were subsequently normalized using Iterative Correction and Eigenvector decomposition (ICE) built in HiC-Pro. FitHiC2 (v2.0.7)^83^ was then used to call chromatin interactions with *-U 2000000 -L 20000 -r 10000 -p 2*. Significant promoter-anchored interactions, declared as chromatin contacts within 1Mb at an *FDR* threshold <1%, were defined based on overlap with gene promoter regions (2kb upstream and downstream of transcription start sites [TSS]). CN Hi-C data was obtained from Hu et al. 2020^17^.

TopDom (v0.9.1)^84^ with default arguments was used to define topologically associating domains (TADs) from normalized 40kb contact matrices. TopDom firstly computed the average contact frequency (defined as a value of *binSignal*) between upstream and downstream regions for each bin. The *binSignal* values demarcate TAD boundaries such that it shows local minimum in a TAD boundary while it is relatively high within a TAD domain. We then used pheatmap (v1.0.12) package to visualize chromatin contact maps.

Information about samples and Hi-C libraries for DN are described below:

| Hi-C library | Sample information | *cis* reads | total reads | *cis* ratio |
| --- | --- | --- | --- | --- |
| 991 | Male, 35 yo<br>African American | 158,222,950 | 230,186,257 | 0.69 |
| 1325 | Female, 45 yo<br>African American | 192,097,648 | 266,025,620 | 0.72 |
| 1423 | Female, 46 yo<br>African American | 164,038,226 | 197,508,199 | 0.83 |
| 1562 | Male, 36 yo<br>Caucasian | 176,575,402 | 237,642,800 | 0.74 |
| 1883 | Male, 45 yo<br>African American | 243,313,838 | 325,461,515 | 0.75 |
| Pooled |  | 934,248,064 | 1,256,824,391 | 0.74 |

### Gene regulatory relationships of cortical and dopaminergic neurons

RNA-seq and ATAC-seq data from CNs and DNs (cortical glutamatergic neurons) were obtained from GEO (GSE129017)^21^. We used FastQC (v.0.11.8)^85^ to check the quality of RNA-seq and ATAC-seq reads.

For RNA-seq analysis, clean reads were mapped to the human reference genome (hg19) from the UCSC database with HISAT2 (v.2.2.1)^86^ using default parameters. We assembled and quantified transcripts using StringTie (v.2.1.2)^87^. Normalized expression values (fragments per kilobase of exon model per million reads mapped, FPKM) of DN marker genes were compared between CNs and DNs.

For ATAC-seq analysis, we applied Bowtie2 (v2.3.5.1)^82^ with *--very-sensitive* to map clean reads from ATAC-seq to hg19 from the UCSC database. After filtering out mitochondrial reads, duplicate reads were further removed by Picard (v.2.20.1, http://broadinstitute.github.io/picard/) by MarkDuplicates function. We then ran MACS2 (v.2.1.0.20150731)^88^ with *--nolambda --nomodel* to call open chromatin regions. Blacklisted regions from ENCODE were removed from the MACS2-called peaks. Finally, we used DiffBind (v.2.13.1)^89^ to analyze differentially open chromatin regions between CN and DN ATAC-seq data. Differentially open chromatin regions were selected on the basis of *FDR<0.05*. Differential ATAC-seq peaks between CNs and DNs were intersected with promoter-anchored interactions to identify enhancer-promoter interactions.

### GWAS datasets

We used the largest publicly available GWAS datasets from European ancestries of cigarette smoking and alcohol use traits. Datasets used were: Problematic alcohol use (PAU)^6^, N = 435,563; Drinks Per Week (DPW)^5^, N = 941,280; Nicotine Dependence (ND)^7^, N = 78,067; Cigarettes Per Day (CPD)^5^, N = 337,334.

### LD score regression analysis

Stratified LD score regression (LDSC)^90^ was used to estimate the enrichment of SNP-based heritability for PAU, DPW, ND, and CPD GWAS. *Cis*-regulatory elements (CREs) of the PFC and substantia nigra (SN) were defined as regions marked as active transcriptional start site (TSSs, state 1), flanking active TSSs (state 2), genic enhancers (state 6) and enhancers (state 7) in the chromHMM core 15-state model^19^. We acquired CREs of CNs and DNs by converting chromatin accessibility peaks reported in Zhang et al^21^ to hg19 using liftOver. CREs of different cell types in the cortex were obtained by merging H3K27ac and H4K3me3 peaks reported from Nott et al^22^. Genetic variants were annotated to corresponding CREs, and SNP-based heritability enrichment was calculated using the GWAS summary statistics mentioned above.

### H-MAGMA and gene selection

H-MAGMA input files were generated from the midbrain DN Hi-C data (DN H-MAGMA). Briefly, exonic and promoter SNPs were assigned to the genes in which they reside, while intronic and intergenic SNPs were coupled to their target genes based on significant chromatin interactions detected in DNs. For CN H-MAGMA, we used H-MAGMA input previously generated from NeuN-positive cells sorted from the dorsolateral prefrontal cortex (DLPFC)^17^. These input files are available in the GitHub repository at https://github.com/thewonlab/H-MAGMA.

Using these input files, we ran Hi-C coupled MAGMA (H-MAGMA) v.1.08^91^as previously described with the following code^12^.

~~~
magma_v1.08/magma -–bfile g1000_eur –pval <GWAS summary statistics>
use=rsid, p ncol=N -–gene-annot <MAGMA input annotation file> -–
out<OUTPUT file>
~~~

H-MAGMA converts SNP-level *p-values* into gene-level *p-values*, from which we selected protein-coding genes that are significantly associated with cigarette smoking and alcohol use traits at *FDR<0.05*. Since we used both cortical and dopaminergic Hi-C datasets, we obtained two gene sets, one from running CN H-MAGMA and the other from running DN H-MAGMA. These genes were used for subsequent functional analyses.

### Gene ontology

We performed gene ontology (GO) analyses to identify biological pathways underlying cigarette smoking and alcohol use traits. Rather than using a selected set of genes with a specific *FDR* cutoff, we ran a rank-based gene ontology analysis using the Bioconductor package g:Profiler (v.0.7.0)^92^. Briefly, genes were ranked based on Z-scores calculated by H-MAGMA, such that genes more significantly associated with a given trait are listed at the top. Biological pathways over-represented by the highly ranked genes were selected.

~~~
gprofiler(<RANKED gene list>, organism=“hsapiens”, ordered_query=T,
significant=T, max_p_value=0.05, min_set_size=15, max_set_size=600,
min_isect_size=5, correction_method=“fdr”, hier_filtering=“strong”,
custom_bg=background gene set, include_graph=T, src_filter=“GO”)
~~~

### Cellular expression

We identified cellular expressions of cigarette smoking and alcohol use risk genes using publicly available single cell RNA sequencing data (scRNA-seq)^50, 54, 62^. Given that GWAS power can influence the number of significant genes for a given trait, we used two different *FDR* thresholds to select cigarette smoking and alcohol use risk genes. We used *FDR<0.1* for GWAS for ND, given that there were <20 genome-wide significant loci; *FDR<0.05* for PAU, DPW, and CPD given >20 genome-wide significant loci. Next, we used scRNA-seq data from the human cortex to annotate cell-type specific and neuronal subcluster specific expressions of CN H-MAGMA risk genes for PAU, DPW, ND, and CPD^50, 62^. Upon gene selection, we scaled expression profiles of each cell using the `scale (x, center=T, scale=F)` function in `R` and calculated the average expression of H-MAGMA risk genes in a given cell. Cell types (e.g. Neurons, Astrocytes, Microglia, Endothelial, Oligodendrocytes) and neuronal subclusters (e.g. excitatory and inhibitory neurons) with highest average expression values were identified as central cell types underlying cigarette smoking and alcohol use traits. Similarly, we annotated DN H-MAGMA risk genes to midbrain cell-types identified from scRNA-seq from the human embryonic ventral midbrain during development^54^. Midbrain cell types include Radial glial (Rgl), Neuroblast, Progenitors (consisting of medial floorplate, lateral floorplate, midline, and basal plate progenitors), Neuronal progenitors (NProg), Oligodendrocyte progenitor cells (OPC), Dopaminergic neurons, Endothelial, GABAergic neurons, Microglial, Oculomotor and trochlear nucleus (OMTN), Pericytes, Red nucleus, and Serotonergic neurons. Lastly, we sought to identify specific dopaminergic clusters enriched for cigarette smoking and alcohol use risk genes. To achieve this, we annotated DN H-MAGMA risk genes to the dopaminergic lineage identified from the human embryonic ventral midbrain^54^.

### Regional expression pattern

We measured brain regional expression profiles of cigarette smoking and alcohol use risk genes using a comprehensive dataset of the mouse nervous system from Zeisel et al. 2018^23^. Of the 24 brain regions represented, we analyzed the following regions: Cortex, Hippocampus, Amygdala, Striatum, Thalamus, Hypothalamus, Midbrain, Cerebellum, and Spinal cord. We generated a new list of risk genes for cigarette smoking and alcohol use traits by combining CN and DN H-MAGMA risk genes using `union(x,y)` in `R` to ensure that our findings were not being dominated by a specific H-MAGMA gene set. Next, we scaled each brain regional expression profile using `scale(x, center=T, scale=F)` in `R` and calculated the average expression of H-MAGMA risk genes. Regions with relatively enriched expression were identified as brain regions associated with cigarette smoking and alcohol use traits.

### Pleiotropic genes

To identify shared neurobiological mechanisms between cigarette smoking and alcohol use traits, we compared gene-level association statistics of PAU and CPD using the rank-rank hypergeometric overlap (RRHO, v.1.40)^65^ R package. Because non-coding genes could result in spurious relationships, we restricted our analysis to protein-coding genes and ran RRHO with the following command line.

~~~
RRHO.result = RRHO (Gene list 1, Gene list 2, outputdir=“∼/output/”,
alternative=“enrichment”, labels=c(“Gene list 1”, “Gene list 2”),
BY=TRUE, log.ind=TRUE, plot=TRUE)
~~~

Overlapping genes between PAU and CPD as identified by RRHO output files served as pleiotropic genes for downstream analyses. To identify biological pathways underlying pleiotropic genes, we ran GO analyses as previously described^92^. Because RRHO does not provide a ranked gene list, we performed GO analyses on the unranked pleiotropic genes with the following command line.

~~~
gprofiler(<UNRANKED pleiotropic gene list>, organism=”hsapiens”,
ordered_query=F, significant=T, max_p_value=0.05, min_set_size=15,
max_set_size=800, min_isect_size=5, correction_method=”fdr”,
hier_filtering=”moderate”, custom_bg=background gene set,
include_graph=T, src_filter=”GO)
~~~

### Differentially expressed genes in response to cocaine

To test for cell-type specific changes of pleiotropic genes in response to cocaine, we first overlapped H-MAGMA genes with differentially expressed genes (DEGs) in the rodent nucleus accumbens (NAc) upon cocaine exposure^66^. DEGs from dopaminergic cell clusters (Drd1-MSNs, Drd2-MSNs1, Drd2-MSNs2, Drd3-MSNs) identified from the NAc were converted from the rodent HUGO Gene Nomenclature Committee (HGNC) symbol to their homologous human Ensembl gene IDs. Rodent genes that did not have a corresponding human Ensembl ID were removed from analysis, resulting in a total of 12,437 cocaine background genes from the dopaminergic clusters. We next selected for DEGs at FDR adjusted *p-value<0.05* from the dataset, resulting in a total of 608 significant dopaminergic DEGs from the cocaine background genes. These 608 significant DEGs were classified as cocaine DEGs for analysis. Because cocaine background genes differ from all H-MAGMA background genes, we generated a comparable H-MAGMA risk gene set by overlapping pleiotropic genes with the cocaine background genes. Next, we compared the proportion of pleiotropic genes that overlapped with cocaine DEGs using the `Venn(x)` function in the `Vennerable` package (v.3.1.0.9000) in `R`. We also applied a Fisher’s exact test to test for significance of overlap as follows:

~~~
fisher.test(matrix(c(overlapping set, Gene list 1-overlapping set, Gene
list 2-overlapping set, cocaine background genes),2,2)
~~~

Lastly, to assess cell-type specific transcriptional changes of pleiotropic genes upon cocaine treatment, we compared transcriptional changes between saline vs. cocaine in the scRNA-seq data^66^. Briefly, we scaled each cell using the `scale(x, center=T, scale=F)`function in `R` and generated box plots comparing saline vs. cocaine treatment for each cluster (e.g. Astrocytes, Dopaminergic neurons, GABAergic neurons, Glutamatergic neurons, Metabotropic glutamate receptor [Grm8-MSN], Microglia, Mural cells, Oligodendrocytes, Polydendrocytes, Interneurons). To test if our risk genes behave differently after cocaine treatment, we compared cellular expression levels between saline and cocaine treatment using the `t.test (x1, x2)` function in `R`.

### Drug enrichment analysis

We used EnrichR^69^ to obtain a list of potential drug candidates for cigarette smoking and alcohol use risk genes. We limited our analysis to the Drug Signature (DsigDB) and Drug Matrix databases of EnrichR as they were the most comprehensive drug libraries available on the platform. Using cigarette smoking and alcohol use risk genes identified with the threshold of *FDR<0.05* for PAU, DPW, ND, and CPD, we adjusted drug associated *p-values* provided by EnrichR after multiple testing correction and selected for significant drugs approved by the Food and Drug Administration (FDA) and small molecules.

### Data Availability

CN (syn21760712) and DN (syn24184521) Hi-C datasets described in this manuscript are available via the PsychENCODE Knowledge Portal (https://psychencode.synapse.org/). The PsychENCODE Knowledge Portal is a platform for accessing data, analyses, and tools generated through grants funded by the National Institute of Mental Health (NIMH) PsychENCODE program. Data is available for general research use according to the following requirements for data access and data attribution: (https://psychencode.synapse.org/DataAccess). H-MAGMA input and output files are available in the Github repository (https://github.com/thewonlab/H-MAGMA). GWAS summary statistics for DPW and CPD were obtained from https://genome.psych.umn.edu/index.php/GSCAN. GWAS summary statistics for ND and PAU were obtained from dbGaP with the accession numbers and phs001532.v1.p1 and phs001672.v3.p1, respectively. RNA-seq and ATAC-seq data from iPSC-derived CNs and DNs were obtained from GSE129017.

### Code Availability

All custom code used in this work is available in the following Github repository: https://github.com/thewonlab/H-MAGMA.

## Acknowledgements

We thank members of the Won lab for helpful discussions and comments about this paper, in particular, Nana Matoba, Won Mah, and Jessica McAfee. We also acknowledge helpful advice and discussion from Jonathan Pollock, Amy Lossie, and Susan Wright. We thank Drs. Stefano Marenco and Barbara Lipska from the Human Brain Collection Core (HBCC, Bethesda, MD) for providing postmortem brain specimens; Mette Peters, Kelsey Montgomery, and Juliane Schneider for assisting data deposition into synapse. This research was supported by the National Institute on Drug Abuse (R21DA051921, H.W., D.B.H., E.O.J.; U01DA048279, S.A), National Institute of Mental Health (R00MH113823, DP2MH122403, H.W.), the NARSAD Young Investigator Award from the Brain and Behavior Research Foundation (H.W.), and the National Science Foundation Graduate Research Fellowship Program (N.Y.A.S.).

## Author Contributions

N.Y.A.S., B.H., D.B.H., E.O.J., S.A., and H.W. designed the research. S.A. supervised DN Hi-C library generation. N.S. and G.B.H sorted dopaminergic nuclei from the midbrain. M.I. generated DN Hi-C libraries. B.H. analyzed ATAC-seq, RNA-seq, and Hi-C data and developed DN H-MAGMA framework. N.Y.A.S. performed H-MAGMA analysis and functional characterization of risk genes. B.C.Q., J.M., E.O.J., and D.B.H. performed meta-analysis of ND GWAS. H.S. performed EnrichR analysis. H.W., N.Y.A.S., and B.H. generated and edited figures. H.W., N.Y.A.S., and B.H. co-wrote the first draft of the manuscript, which was subsequently revised by all other co-authors.

## Supplementary Table Legend

**Supplementary Table 1**

Sample metadata description for dopaminergic neuronal Hi-C data.

**Supplementary Table 2**

H-MAGMA output files for cigarette smoking and alcohol use traits based on cortical and dopaminergic neuronal Hi-C data.

**Supplementary Table 3**

Risk genes identified by cortical and dopaminergic H-MAGMA.

**Supplementary Table 4**

Gene ontology output files for cigarette smoking and alcohol use traits.

**Supplementary Table 5**

List of pleiotropic genes between problematic alcohol use and cigarettes per day.

**Supplementary Table 6**

Drug enrichment analysis of cigarette smoking and alcohol use traits.

## Supplementary Figure Legend

**Supplementary Figure 1.**
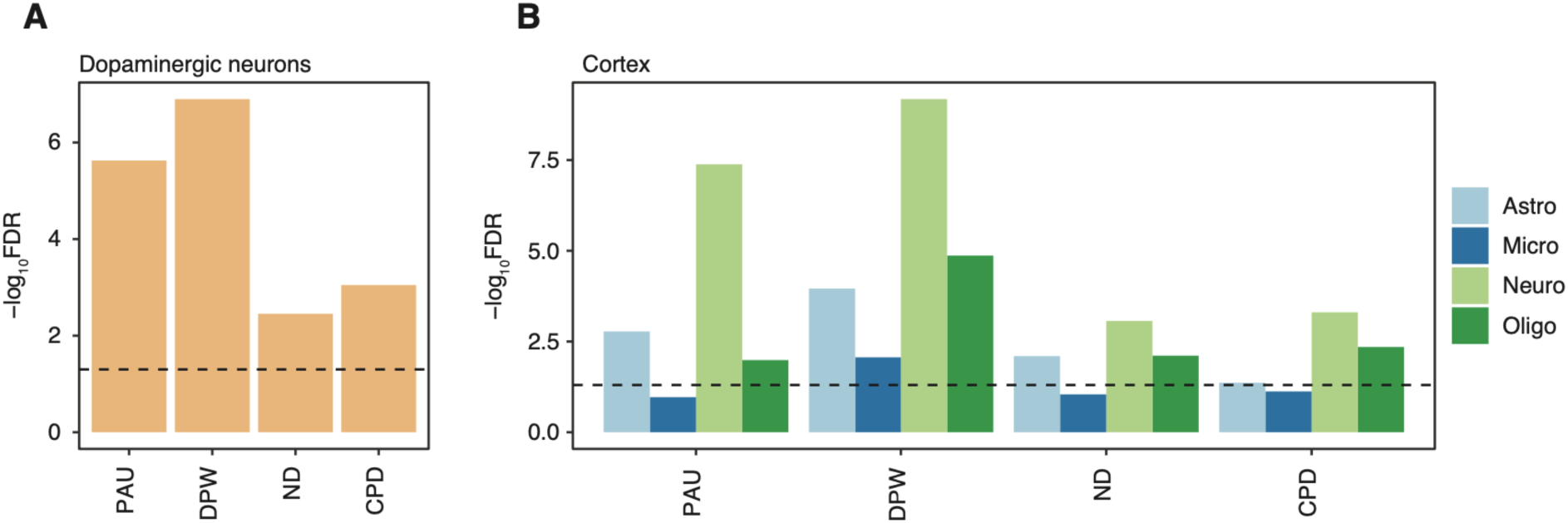
Heritability enrichment of cigarette smoking and alcohol use traits in DNs and cortical cell types. **A**. Heritability enrichment of cigarette smoking and alcohol use traits using stratified LDSC. Genetic risk variants associated with cigarette smoking and alcohol use traits are enriched for DN-CREs. **B**. Cell-type specific heritability enrichment of cigarette smoking and alcohol use traits in the cortex. We observed neuronal enrichment for cigarette smoking and alcohol use traits. Dotted line indicates *FDR=0.05*. Astro, astrocyte; Micro, microglia; Neuro, neuron; Oligo, oligodendrocyte.

**Supplementary Figure 2.**
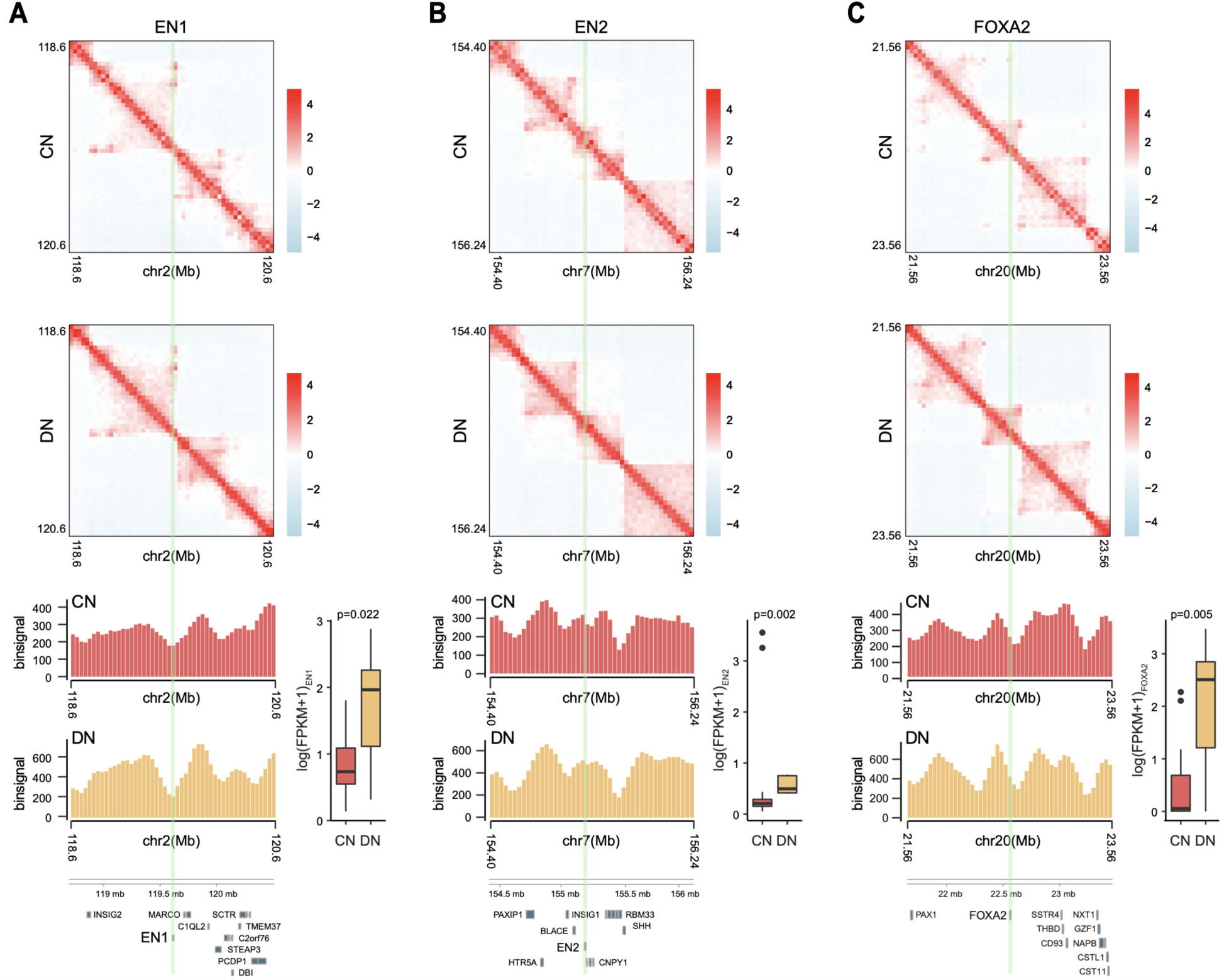
Difference in chromatin architecture between CNs and DNs. Normalized contact frequency matrices for *EN1* (**A**), *EN2* (**B**), and *FOXA2* (**C**) loci (highlighted in green). Heatmaps represent 40kb normalized Hi-C contact matrices of CNs (top) and DNs (middle). Z-axis, normalized contact frequency. Barplots in the bottom represent *binSingal* values calculated from normalized Hi-C contact matrices of CNs (red) and DNs (orange). Normalized expression values of *EN1*, *EN2*, and *FOXA2* demonstrate their elevated expression in DN. Wilcoxon Rank Sum test was used to calculate *p* values.

**Supplementary Figure 3.**
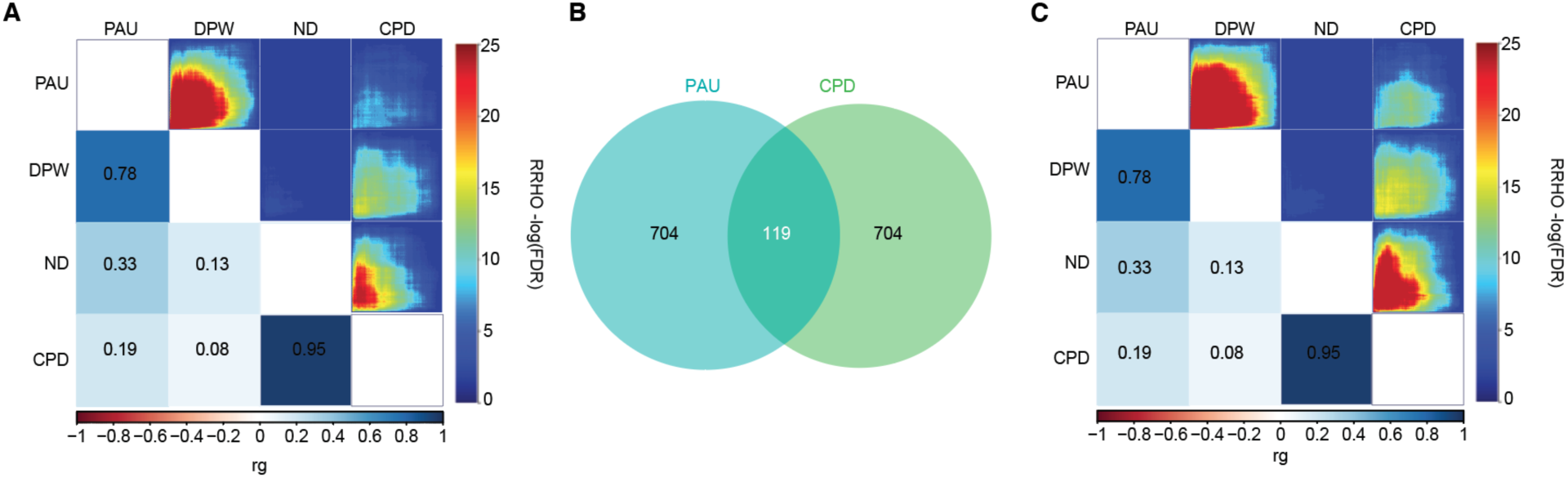
Genetic correlations and overlapping genes between cigarette smoking and alcohol use traits. **A**. LDSC and RRHO were used to estimate genetic correlations and gene-level overlap between cigarette smoking and alcohol use traits, respectively. Bottom left plot represents genetic correlations (rg) while top right plot denotes gene-level overlap using CN H-MAGMA output files. **B**. Overlap between PAU and CPD risk genes identified by CN H-MAGMA using RRHO. **C**. Genetic correlations (bottom left) and gene-level overlap from DN H-MAGMA output files.

**Supplementary Figure 4.**
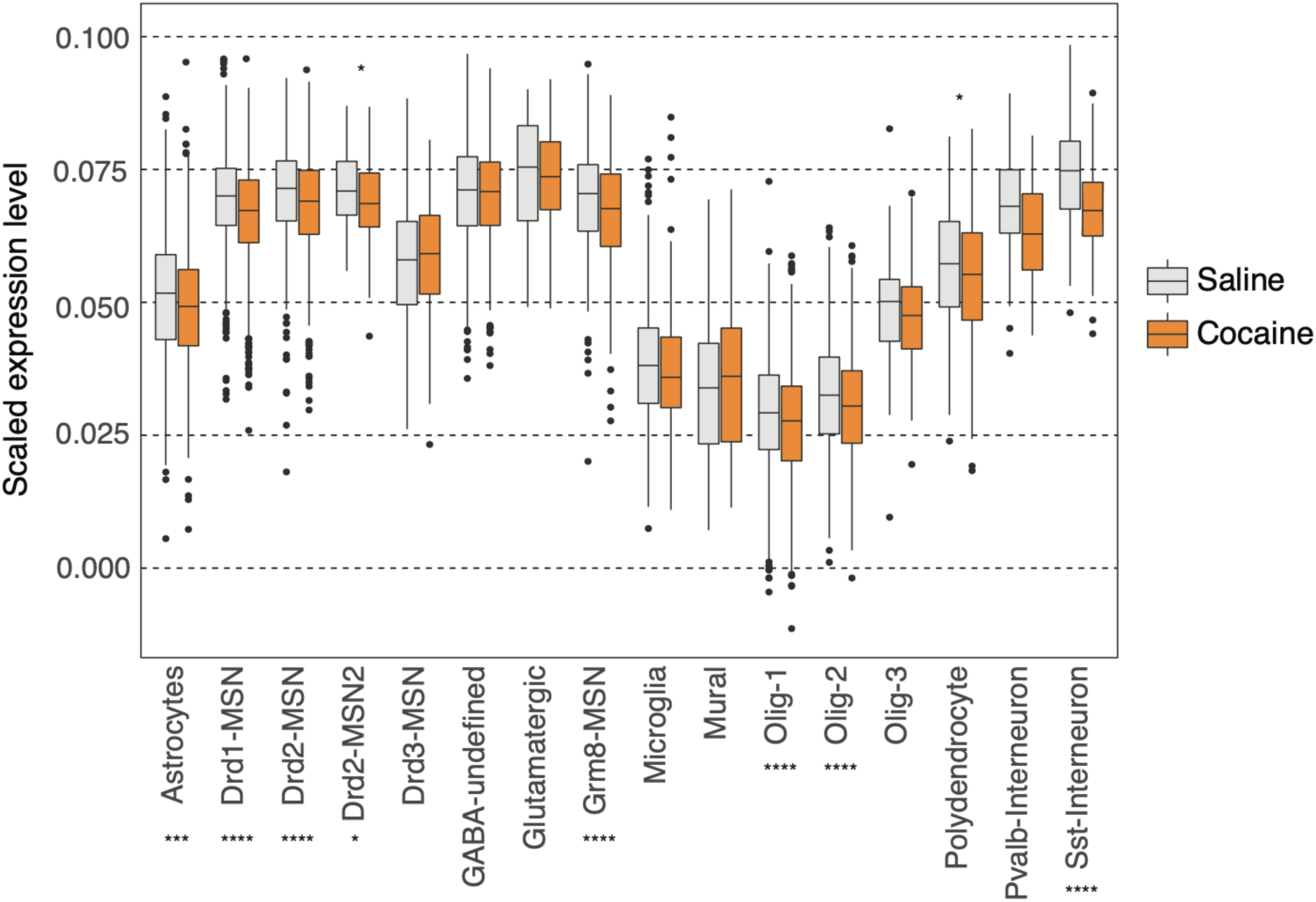
Cellular expression changes of pleiotropic genes in response to cocaine treatment. The x-axis indicates cell types identified from the mouse NAc while the y-axis indicates scaled expression values of pleiotropic genes within a given cluster. T-tests were used to measure significance. FDR of significant pairs are represented with asterisks. * FDR adjusted p*<0.05*; *** FDR adjusted p*<0.001*; **** FDR adjusted p*<0.0001*. Drd1-MSN, dopamine receptor D1 medium spiny neurons enriched for D1-like family; Drd2-MSN and Drd2-MSN2, dopamine receptor D2 medium spiny neurons; Drd3-MSN, dopamine receptor D3 medium spiny neurons; Grm8-MSN, glutamatergic neurons metabotropic glutamate receptor; Oligo, oligodendrocyte.

## Reference

1. Substance Abuse and Mental Health Services Administration. (2019). Key substance use and mental health indicators in the United States: Results from the 2018 National Survey on Drug Use and Health (HHS Publication No. PEP19-5068, NSDUH Series H-54). Rockville, MD: Center for Behavioral Health Statistics and Quality, Substance Abuse and Mental Health Services Administration.

2. Peacock, A. et al. Global statistics on alcohol, tobacco and illicit drug use: 2017 status report. Addiction 113, 1905–1926 (2018).

3. National Center for Chronic Disease Prevention and Health Promotion (US) Office on Smoking and Health. The Health Consequences of Smoking—50 Years of Progress: A Report of the Surgeon General. (Centers for Disease Control and Prevention (US), 2014).

4. Kranzler, H. R. et al. Genome-wide association study of alcohol consumption and use disorder in 274,424 individuals from multiple populations. Nat. Commun. 10, 1499 (2019).

5. Liu, M. et al. Association studies of up to 1.2 million individuals yield new insights into the genetic etiology of tobacco and alcohol use. Nat. Genet. 51, 237–244 (2019).

6. Zhou, H. et al. Genome-wide meta-analysis of problematic alcohol use in 435,563 individuals yields insights into biology and relationships with other traits. Nat. Neurosci. 23, 809–818 (2020).

7. Quach, B. C. et al. Expanding the genetic architecture of nicotine dependence and its shared genetics with multiple traits. Nat. Commun. 11, 1–13 (2020).

8. Dekker, J. Gene regulation in the third dimension. Science 319, 1793–1794 (2008).

9. Won, H. et al. Chromosome conformation elucidates regulatory relationships in developing human brain. Nature 538, 523–527 (2016).

10. Sanyal, A., Lajoie, B. R., Jain, G. & Dekker, J. The long-range interaction landscape of gene promoters. Nature 489, 109–113 (2012).

11. Mah, W. & Won, H. The three-dimensional landscape of the genome in human brain tissue unveils regulatory mechanisms leading to schizophrenia risk. Schizophr. Res. 217, 17–25 (2020).

12. Sey, N. Y. A. et al. A computational tool (H-MAGMA) for improved prediction of brain-disorder risk genes by incorporating brain chromatin interaction profiles. Nat. Neurosci. (2020) doi:10.1038/s41593-020-0603-0.

13. Rajarajan, P. et al. Neuron-specific Signatures in the Chromosomal Connectome Are Associated with Schizophrenia Risk. Science Accepted f, eaat4311–eaat4311 (2018).

14. Koob, G. F. & Volkow, N. D. Neurobiology of addiction: a neurocircuitry analysis. Lancet Psychiatry 3, 760–773 (2016).

15. Lammel, S., Lim, B. K. & Malenka, R. C. Reward and aversion in a heterogeneous midbrain dopamine system. Neuropharmacology 76 Pt B, 351–359 (2014).

16. Clarke, R. & Adermark, L. Dopaminergic Regulation of Striatal Interneurons in Reward and Addiction: Focus on Alcohol. Neural Plast. 2015, 814567 (2015).

17. Hu, B. et al. Neuronal and glial 3D chromatin architecture illustrates cellular etiology of brain disorders. 2020.05.14.096917 (2020) doi:10.1101/2020.05.14.096917.

18. Espeso-Gil, S. et al. A chromosomal connectome for psychiatric and metabolic risk variants in adult dopaminergic neurons. Genome Med. 12, 19 (2020).

19. Consortium, Roadmap Epigenomics et al. Integrative analysis of 111 reference human epigenomes. Nature 518, 317–330 (2015).

20. Berke, J. D. & Hyman, S. E. Addiction, dopamine, and the molecular mechanisms of memory. Neuron 25, 515–532 (2000).

21. Zhang, S. et al. Allele-specific open chromatin in human iPSC neurons elucidates functional disease variants. Science vol. 369 561–565 (2020).

22. Nott, A. et al. Brain cell type-specific enhancer-promoter interactome maps and disease-risk association. Science 366, 1134–1139 (2019).

23. Zeisel, A. et al. Molecular Architecture of the Mouse Nervous System. Cell 174, 999– 1014.e22 (2018).

24. Metzakopian, E. et al. Genome-wide characterization of Foxa2 targets reveals upregulation of floor plate genes and repression of ventrolateral genes in midbrain dopaminergic progenitors. Development 139, 2625–2634 (2012).

25. . Lee, H.-S., et al. Foxa2 and Nurr1 synergistically yield A9 nigral dopamine neurons exhibiting improved differentiation, function, and cell survival. Stem Cells 28, 501–512 (2010).

26. Saucedo-Cardenas, O. et al. Nurr1 is essential for the induction of the dopaminergic phenotype and the survival of ventral mesencephalic late dopaminergic precursor neurons. Proc. Natl. Acad. Sci. U. S. A. 95, 4013–4018 (1998).

27. Dixon, J. R. et al. Topological domains in mammalian genomes identified by analysis of chromatin interactions. Nature 485, 376–380 (2012).

28. Simon, H. H., Saueressig, H., Wurst, W., Goulding, M. D. & O’Leary, D. D. Fate of midbrain dopaminergic neurons controlled by the engrailed genes. J. Neurosci. 21, 3126–3134 (2001).

29. Delatour, L. C., Yeh, P. W. & Yeh, H. H. Ethanol Exposure In Utero Disrupts Radial Migration and Pyramidal Cell Development in the Somatosensory Cortex. Cereb. Cortex 29, 2125–2139 (2019).

30. Skorput, A. G. J., Gupta, V. P., Yeh, P. W. L. & Yeh, H. H. Persistent Interneuronopathy in the Prefrontal Cortex of Young Adult Offspring Exposed to Ethanol In Utero. J. Neurosci. 35, 10977–10988 (2015).

31. Kazemi, T. et al. Investigating the influence of perinatal nicotine and alcohol exposure on the genetic profiles of dopaminergic neurons in the VTA using miRNA–mRNA analysis. Scientific Reports vol. 10 (2020).

32. Linker, K. E., Cross, S. J. & Leslie, F. M. Glial mechanisms underlying substance use disorders. Eur. J. Neurosci. 50, 2574–2589 (2019).

33. Lacagnina, M. J., Rivera, P. D. & Bilbo, S. D. Glial and Neuroimmune Mechanisms as Critical Modulators of Drug Use and Abuse. Neuropsychopharmacology 42, 156–177 (2017).

34. Fox, H. C., Milivojevic, V., Angarita, G. A., Stowe, R. & Sinha, R. Peripheral immune system suppression in early abstinent alcohol-dependent individuals: Links to stress and cue-related craving. J. Psychopharmacol. 31, 883–892 (2017).

35. Zuluaga, P. et al. Wide array of T-cell subpopulation alterations in patients with alcohol use disorders. Drug Alcohol Depend. 162, 124–129 (2016).

36. Milivojevic, V. et al. Peripheral Immune System Adaptations and Motivation for Alcohol in Non-Dependent Problem Drinkers. Alcoholism: Clinical and Experimental Research vol. 41 585–595 (2017).

37. Pasala, S., Barr, T. & Messaoudi, I. Impact of alcohol abuse on the adaptive immune system. Alcohol Res. 37, 185 (2015).

38. Higuchi, T. et al. Current cigarette smoking is a reversible cause of elevated white blood cell count: Cross-sectional and longitudinal studies. Prev Med Rep 4, 417–422 (2016).

39. Díaz-Villanueva, J. F., Díaz-Molina, R. & García-González, V. Protein Folding and Mechanisms of Proteostasis Int. J. Mol. Sci. 16, 17193–17230 (2015).

40. Topiwala, A. & Ebmeier, K. P. Effects of drinking on late-life brain and cognition. Evid. Based. Ment. Health 21, 12–15 (2018).

41. Kalapatapu, R. K. et al. Substance use history in behavioral-variant frontotemporal dementia versus primary progressive aphasia. J. Addict. Dis. 35, 36–41 (2016).

42. Elman, I. & Borsook, D. Common Brain Mechanisms of Chronic Pain and Addiction. Neuron 89, 11–36 (2016).

43. Maleki, N., Tahaney, K., Thompson, B. L. & Oscar-Berman, M. At the intersection of alcohol use disorder and chronic pain. Neuropsychology 33, 795–807 (2019).

44. Goodman, J. & Packard, M. G. Memory Systems and the Addicted Brain. Front. Psychiatry 7, 24 (2016).

45. Morin, J.-F. G. et al. A Population-Based Analysis of the Relationship Between Substance Use and Adolescent Cognitive Development. Am. J. Psychiatry 176, 98–106 (2019).

46. Wilkinson, A. N. et al. Effects of binge alcohol consumption on sleep and inflammation in healthy volunteers. J. Int. Med. Res. 46, 3938–3947 (2018).

47. Elmenhorst, E.-M. et al. Cognitive impairments by alcohol and sleep deprivation indicate trait characteristics and a potential role for adenosine A1 receptors. Proc. Natl. Acad. Sci. U. S. A. 115, 8009–8014 (2018).

48. Saba, R., Halytskyy, O., Saleem, N. & Oliff, I. A. Buccal Epithelium, Cigarette Smoking, and Lung Cancer: Review of the Literature. Oncology 93, 347–353 (2017).

49. Xu, Z. et al. Cancer mortality attributable to cigarette smoking in 2005, 2010 and 2015 in Qingdao, China. PLoS One 13, e0204221 (2018).

50. Darmanis, S. et al. A survey of human brain transcriptome diversity at the single cell level. Proc. Natl. Acad. Sci. U. S. A. 112, 7285–7290 (2015).

51. Gerfen, C. R., Economo, M. N. & Chandrashekar, J. Long distance projections of cortical pyramidal neurons. J. Neurosci. Res. 96, 1467–1475 (2018).

52. Kim, E. J., Juavinett, A. L., Kyubwa, E. M., Jacobs, M. W. & Callaway, E. M. Three Types of Cortical Layer 5 Neurons That Differ in Brain-wide Connectivity and Function. Neuron 88, 1253–1267 (2015).

53. Matzeu, A. & Martin-Fardon, R. Drug Seeking and Relapse: New Evidence of a Role for Orexin and Dynorphin Co-transmission in the Paraventricular Nucleus of the Thalamus. Front. Neurol. 9, 720 (2018).

54. La Manno, G. et al. Molecular Diversity of Midbrain Development in Mouse, Human, and Stem Cells. Cell 167, 566–580.e19 (2016).

55. Tritsch, N. X., Oh, W.-J., Gu, C. & Sabatini, B. L. Midbrain dopamine neurons sustain inhibitory transmission using plasma membrane uptake of GABA, not synthesis. Elife 3, e01936 (2014).

56. Morello, F. & Partanen, J. Diversity and development of local inhibitory and excitatory neurons associated with dopaminergic nuclei. FEBS Lett. 589, 3693–3701 (2015).

57. Kirby, L. G., Zeeb, F. D. & Winstanley, C. A. Contributions of serotonin in addiction vulnerability. Neuropharmacology 61, 421–432 (2011).

58. Gould, T. J. Nicotine and hippocampus-dependent learning. Mol. Neurobiol. 34, 93–107 (2006).

59. Kutlu, M. G. & Gould, T. J. Nicotinic receptors, memory, and hippocampus. Curr. Top. Behav. Neurosci. 23, 137–163 (2015).

60. Zhu, Y., Wienecke, C. F. R., Nachtrab, G. & Chen, X. A thalamic input to the nucleus accumbens mediates opiate dependence. Nature 530, 219–222 (2016).

61. Chye, Y. et al. Subcortical surface morphometry in substance dependence: An ENIGMA addiction working group study. Addict. Biol. 25, e12830 (2020).

62. Lake, B. B. et al. Neuronal subtypes and diversity revealed by single-nucleus RNA sequencing of the human brain. Science 352, 1586–1590 (2016).

63. Abuse, S. Mental Health Services Administration. (2018). Key substance use and mental health indicators in the United States: Results from the 2017 National Survey on Drug Use and Health (HHS Publication No. SMA 18-5068, NSDUH Series H-53). Rockville, MD: Center for Behavioral Health Statistics and Quality. Substance Abuse and Mental Health Services Administration. Retrieved from https://www.samhsa.gov/data (2019).

64. Bulik-Sullivan, B. et al. An atlas of genetic correlations across human diseases and traits. Nat. Genet. 47, 1236–1241 (2015).

65. Plaisier, S. B., Taschereau, R., Wong, J. A. & Graeber, T. G. Rank-rank hypergeometric overlap: identification of statistically significant overlap between gene-expression signatures. Nucleic Acids Res. 38, e169 (2010).

66. Savell, K. E. et al. A dopamine-induced gene expression signature regulates neuronal function and cocaine response. Sci Adv 6, eaba4221 (2020).

67. Hendershot, C. S., Wardell, J. D., Samokhvalov, A. V. & Rehm, J. Effects of naltrexone on alcohol self-administration and craving: meta-analysis of human laboratory studies. Addiction Biology vol. 22 1515–1527 (2017).

68. Stead, L. F. et al. Nicotine replacement therapy for smoking cessation. Cochrane Database Syst. Rev. 11, CD000146 (2012).

69. Kuleshov, M. V. et al. Enrichr: a comprehensive gene set enrichment analysis web server 2016 update. Nucleic Acids Res. 44, W90–7 (2016).

70. Castillo-Carniglia, A., Keyes, K. M., Hasin, D. S. & Cerdá, M. Psychiatric comorbidities in alcohol use disorder. Lancet Psychiatry 6, 1068–1080 (2019).

71. Murthy, P., Mahadevan, J. & Chand, P. K. Treatment of substance use disorders with co-occurring severe mental health disorders. Curr. Opin. Psychiatry 32, 293–299 (2019).

72. Hartz, S. M. et al. Association Between Substance Use Disorder and Polygenic Liability to Schizophrenia. Biol. Psychiatry 82, 709–715 (2017).

73. Chang, L.-H. et al. Associations between polygenic risk for tobacco and alcohol use and liability to tobacco and alcohol use, and psychiatric disorders in an independent sample of 13,999 Australian adults. Drug and Alcohol Dependence vol. 205 107704 (2019).

74. Hoffman, J. L. et al. Alcohol drinking exacerbates neural and behavioral pathology in the 3xTg-AD mouse model of Alzheimer’s disease. Int. Rev. Neurobiol. 148, 169–230 (2019).

75. Nicholatos, J. W. et al. Nicotine promotes neuron survival and partially protects from Parkinson’s disease by suppressing SIRT6. Acta Neuropathologica Communications vol. 6 (2018).

76. Piao, W.-H. et al. Nicotine and inflammatory neurological disorders. Acta Pharmacologica Sinica vol. 30 715–722 (2009).

77. Bush, T., Lovejoy, J. C., Deprey, M. & Carpenter, K. M. The effect of tobacco cessation on weight gain, obesity, and diabetes risk. Obesity vol. 24 1834–1841 (2016).

78. Germeroth, L. J. & Levine, M. D. Postcessation weight gain concern as a barrier to smoking cessation: Assessment considerations and future directions. Addict. Behav. 76, 250–257 (2018).

79. McCrory, E. J. & Mayes, L. Understanding Addiction as a Developmental Disorder: An Argument for a Developmentally Informed Multilevel Approach. Current Addiction Reports vol. 2 326–330 (2015).

80. Semick, S. A. et al. Developmental effects of maternal smoking during pregnancy on the human frontal cortex transcriptome. Mol. Psychiatry 25, 3267–3277 (2018).

81. Servant, N. et al. HiC-Pro: an optimized and flexible pipeline for Hi-C data processing. Genome Biol. 16, 259 (2015).

82. Langmead, B. & Salzberg, S. L. Fast gapped-read alignment with Bowtie 2. Nat. Methods 9, 357–359 (2012).

83. Kaul, A., Bhattacharyya, S. & Ay, F. Identifying statistically significant chromatin contacts from Hi-C data with FitHiC2. Nat. Protoc. 15, 991–1012 (2020).

84. Shin, H. et al. TopDom: an efficient and deterministic method for identifying topological domains in genomes. Nucleic Acids Res. 44, e70 (2016).

85. Andrews, S. & Others. FastQC: a quality control tool for high throughput sequence data. (2010).

86. Kim, D., Paggi, J. M., Park, C., Bennett, C. & Salzberg, S. L. Graph-based genome alignment and genotyping with HISAT2 and HISAT-genotype. Nat. Biotechnol. 37, 907–915 (2019).

87. Pertea, M. et al. StringTie enables improved reconstruction of a transcriptome from RNA-seq reads. Nat. Biotechnol. 33, 290–295 (2015).

88. Zhang, Y. et al. Model-based analysis of ChIP-Seq (MACS). Genome Biol. 9, R137 (2008).

89. Stark, R., Brown, G. & Others. DiffBind: differential binding analysis of ChIP-Seq peak data. R package version 100, 4–3 (2011).

90. Finucane, H. K. et al. Partitioning heritability by functional annotation using genome-wide association summary statistics. Nat. Genet. 47, 1228–1235 (2015).

91. de Leeuw, C., Sey, N. Y. A., Posthuma, D. & Won, H. A response to Yurko et al: H-MAGMA, inheriting a shaky statistical foundation, yields excess false positives. 2020.09.25.310722 (2020) doi:10.1101/2020.09.25.310722.

92. Reimand, J., Kull, M., Peterson, H., Hansen, J. & Vilo, J. g:Profiler--a web-based toolset for functional profiling of gene lists from large-scale experiments. Nucleic Acids Res. 35, W193–200 (2007).

